# PGC-1α coordinates with Bcl-2 to control cell cycle in U251 cells through reducing ROS

**DOI:** 10.1101/111567

**Authors:** Kun Yao, Xufeng Fu, Du Xing, Yan Li, Bing Han, Chen Zexi, Shanshan Yang, Ran Wei, Jiaqi Zhou, Qinghua Cui

## Abstract

B-cell lymphoma 2 (*Bcl-2*) has a dual function, acting both as an oncogene and an anti-tumor gene. It is well known that Bcl-2 exerts its tumor promoting function through the mitochondrial pathway. However, the mechanism by which Bcl-2 suppresses tumor formation is not well understood. We have previously shown that Bcl-2 inhibits cell cycle progression from the G_0_/G_1_ to the S phase after serum starvation, and that quiescent Bcl-2 expressing cells maintained a significant lower level of mitochondrial reactive oxygen species (ROS) than the control cells. Based on the fact that ROS mediate cell cycle progression, and are controlled by peroxisome proliferator-activated receptor-γ co-activator 1α (PGC-1α), a key molecule induced by prolonged starvation and involved in mitochondrial metabolism, we hypothesized that PGC-1α might be related with the cell cycle function of Bcl-2. Here, we showed that PGC-1α was upregulated upon Bcl-2 overexpression and downregulated following Bcl-2 knockdown during serum starvation. Knockdown of PGC-1α activated Bcl-2 expression. Taken together, our results suggest that after serum depletion, PGC-la might coordinate with Bcl-2 to reduce ROS, which in turn delay cell cycle progression.

**Summary statement:** PGC-1α coordinate with Bcl-2 delay cell cycle progression to reduce ROS after serum depletion in human glioma U251 cells.

## Introduction

B-cell lymphoma 2 (*Bcl-2*) has both pro-apoptotic and anti-apoptotic potentials. It is well known that Bcl-2 protein is anchored to the mitochondrial outer membrane, and antagonizes with the pro-apoptotic protein BAX by forming Bcl-2/BAX heterodimers that control mitochondrial membrane permeability and promote tumorigenesis (Hockenbery et al., 1990). Bcl-2 acts as an anti-tumor gene in early stage solid tumors, where it significantly reduces tumor incidence by regulating the cell cycle (Vail et al., 2001; Murphy et al., 1999). *In vitro* cell studies demonstrated that the most pronounced cell cycle effect of Bcl-2 is delay of progression to S phase from G0/G1 (Janumyan et al., 2008;Janumyan et al., 2003), which has also been confirmed in a Bcl-2 transgenic mouse model, where proliferation of lymphoid T cells was restrained and tumor-associated morbidity was significantly decreased (Cheng et al., 2004). However, the signaling pathways by which Bcl-2 suppresses tumor formation are not fully understood.

We have recently observed that quiescent Bcl-2 overexpressing cells maintain less mitochondrial ROS (Reactive Oxygen Species) than control cells, elevation of ROS accelerates cell cycle progression (unpublished data). These findings strongly indicated that mitochondrial oxidative phosphorylation (OXPHOS) plays a key role in the anti-proliferation function of Bcl-2. The most important factor identified to modulate mitochondrial biogenesis and bioenergetics is peroxisome proliferator-activated receptor-γ co-activator 1α (PGC-1α), which is predominantly expressed in tissues with high energy demands, such as the heart, brain, and muscle (McBride et al., 2006;Renee et al., 2008; Esterbauer et al., 1999). As a co-activator, PGC-1α interacts with a broad range of transcription factors and plays an important role in glucose metabolism and prolonged starvation (Meirhaeghe et al., 2003; Lin et al., 2005). PGC-1α can translocate from the nucleus into the mitochondrial to modulate mitochondrial function (Aquilano et al., 2010; Safdar et al., 2011; Smith et al., 2013). Furthermore, our recent findings uncovered that PGC-1α regulates the cell cycle through ATP and ROS signals in fibroblast cells (Fu et al., 2016). Therefore, we proposed that PGC-1α might translocate into mitochondria to coordinate with Bcl-2’s tumor suppression functions through reducing ROS.

In this study, we investigated the relationship between Bcl-2 and PGC-1α. We first establish Bcl-2 stable overexpressing cells and the control cells, synchronize them in G_0_/G_1_ phase using serum starvation (SS) or contact inhibition (CI), then determine the effects of Bcl-2 over-expression or silencing on PGC-1α expression. Similarly, Bcl-2 expression were detected in cells expressing high levels of PGC-1α or after knockdown of PGC-1α. We found that PGC-1α positively correlates with Bcl-2, Bcl-2 negatively responds to PGC-1α expression.

## Results

### PGC-1α expression increased in cells overexpressing Bcl-2

Mouse embryonic fibroblast NIH3T3 cells were transfected with *Bcl-2* cloned in a pBABEpuro plasmid (referred to as 3T3Bcl-2 in this paper). Cells transfected with the empty vector served as the control (referred to as 3T3PB in this paper). We have reported that PGC-1α inhibits the cell cycle progress, and Bcl-2 exerts its anti-tumor function by delay cell cycle progress only at the G_0_/G_1_ stage (Fu et al., 2016; Du et al., 2017; Janumyan et al., 2008). Therefore, here, we first synchronize cells at the G0/G1 stage by serum starvation, then compare PGC-1α expression in 3T3Bcl-2 cells with 3T3PB cells. Our cell cycle profiles showed that both 3T3PB and 3T3Bcl-2 were arrested successfully in G0/G1 phase, the proportion of cells in S phase dropped from ~20% (normal growing, NG) to less than 3% (serum starved, SS) for both cells (Fig. 1A and 1B). We also observed a significant elevation in p27 levels in SS3T3Bcl-2 cells, confirming that Bcl-2 function as tumor-repressive gene through upregulating p27 (Fig. 1C and 1D). At this stage, PGC-1α expression was clearly increased in SS3T3Bcl-2 comparing with NG3T3Bcl-2 cells, while there was no significant difference between control cells SS3T3PB and NG3T3PB (Fig. 1C and 1E). This result suggests that PGC-1α expression associates with Bcl-2 after serum depletion.

**Fig. 1.**
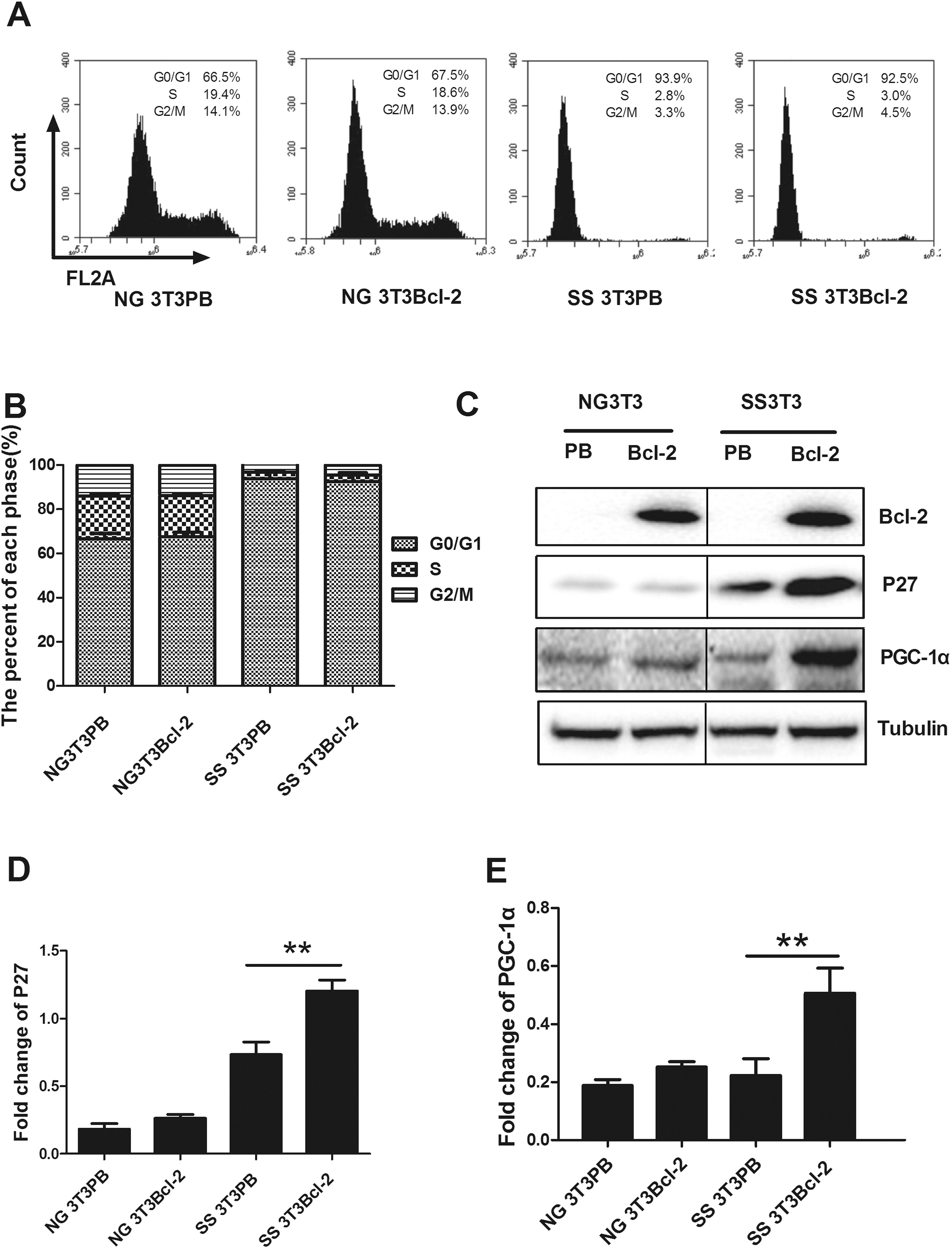
Comparison of p27 and PGC-lα expression between SS-treated and NG 3T3 cells. (A) 3T3PB (pBABEpuro, empty vector) or 3T3Bcl-2 (pBABpuro-Bcl-2) cells were cultured for 48 h in 0.1% serum and harvested for cell cycle analysis by flow cytometry. (B) The percentage of SS-treated or NG 3T3 cells in each cell cycle phase is shown. (C) Expression of PGC-1α and p27 in 3T3 cells was measured by western blot. (D) and (E) Quantification of PGC-1α and p27 protein expression shown in (C), ** denotes p < 0.01, * denotes p < 0.05 (n = 3).

To avoid experimental method bias, the results were verified in cells arrested by contact inhibition (CI). Consistent with the results from the SS method (Fig. 1), both 3T3PB and 3T3Bcl-2 cells were arrested in G_0_/G_1_ phase (CI3T3PB and CI3T3Bcl-2, respectively) (Fig. 2A and 2B), PGC-1α and p27 expression increased significantly in CI3T3Bcl-2 compared to either NG3T3Bcl-2 or CI3T3PB cells, whereas, PGC-1α expression did not significantly change between 3T3PB and CI3T3PB cells (Fig. 2C-Fig. 2E). Based on the combined results from SS (Fig. 1) and CI (Fig. 2) treatment, we conclude that during serum starvation, Bcl-2 delay cell cycle progress and upregulates PGC-1α expression.

**Fig. 2.**
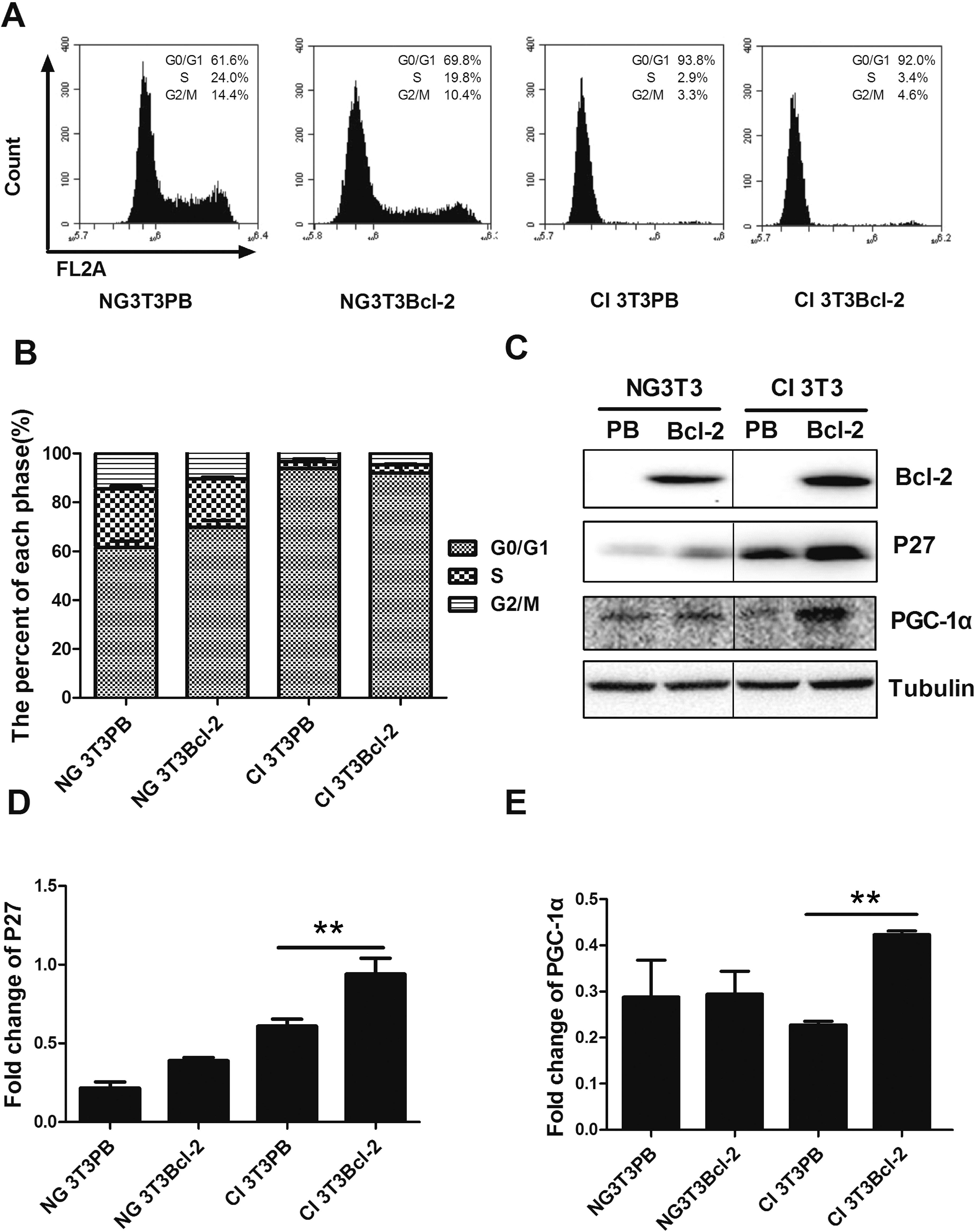
Comparison of p27 and PGC-1α expression between CI-treated and NG 3T3 cells. (A) 3T3PB (pBABEpuro, empty vector) or 3T3Bcl-2 (pBABpuro-Bcl-2) cells were contact inhibited for 72 h and harvested for cell cycle analysis by flow cytometry. (B) The percentage of CI-treated or NG 3T3 cells in each cell cycle phase is shown. (C) Expression of PGC-1α and p27 in 3T3 cells was measured by western blot. (D) and (E) Quantification of PGC-1α and p27 proteins expression shown in (C), ** denotes p < 0.01, * denotes p < 0.05 (n = 3).

### PGC-1α decreased after Bcl-2 knockdown

Since PGC-1α was upregulated in Bcl-2 overexpressing cells (Fig. 1, 2), we further investigated the effects of Bcl-2 knockdown on PGC-1α using glioma U251 cells, which have high Bcl-2 and PGC-1α endogenous expression. Three specific siRNAs (506, 528, and 928) were used to interfere with Bcl-2 expression in U251 cells, while a GAPDH siRNA and a mimic siRNA were used as a positive control and negative control (NC), respectively (Fig. 3A). Western blot confirmed successful Bcl-2 knockdown by all three specific siRNAs, without interfere on positive or negative controls (Fig. 3B). PGC-1α expression was decreased significantly in cells transfected with siRNAs fragment 528 and 928 than 526 which showing less interfering efficiency, PGC-1α expression had no change on controls (Fig. 3C). These results demonstrated that PGC-1α expression was downregulated by Bcl-2 in a dose dependent manner, suggesting the close link between PGC-1α and Bcl-2.

**Fig. 3.**
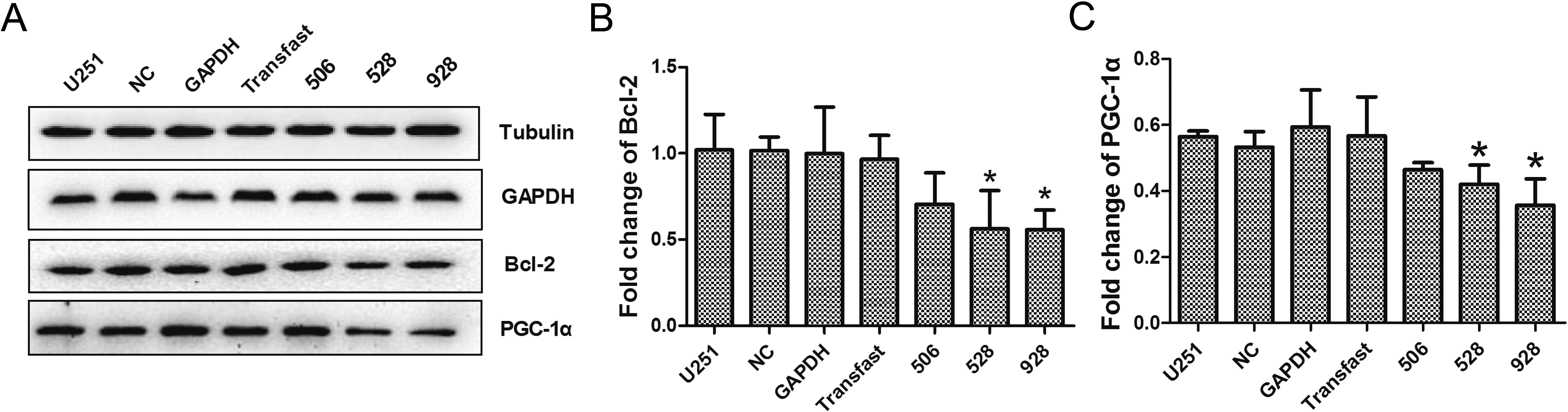
Bcl-2 and PGC-1α knockdown by siRNA in U251 cell. (A) GAPDH, Bcl-2 and PGC-1α expression was detected by western blot after transfection with Bcl-2 specific siRNAs (506, 528 and 928). (B) and (C) Quantification of Bcl-2 and PGC-1α protein expression shown in (A). NC: random siRNA serving as a negative control, GAPDH: positive control, Transfast: transfection reagents only.

### Bcl-2 upregulation correlates with PGC-1α downregulation in U251 cells after SS treatment

Next, we investigated whether Bcl-2 was regulated by PGC-1a. Human glioma cell line U251 has high endogenous expression of PGC-1α and Bcl-2, so we chose these cells to knockdown PGC-1α and check the effects on Bcl-2. U251 cells were treated by SS for 24, 48, and 72 h, and normal growing (NGU251) cells were used as a control. The results showed that during SS treatment, S phase cells dropped down (Fig. 4A and 4B), Bcl-2 was upregulated following PGC-1α downregulation (Fig. 4C). These results suggested that Bcl-2 responds to PGC-1α in a negative way.

**Fig. 4.**
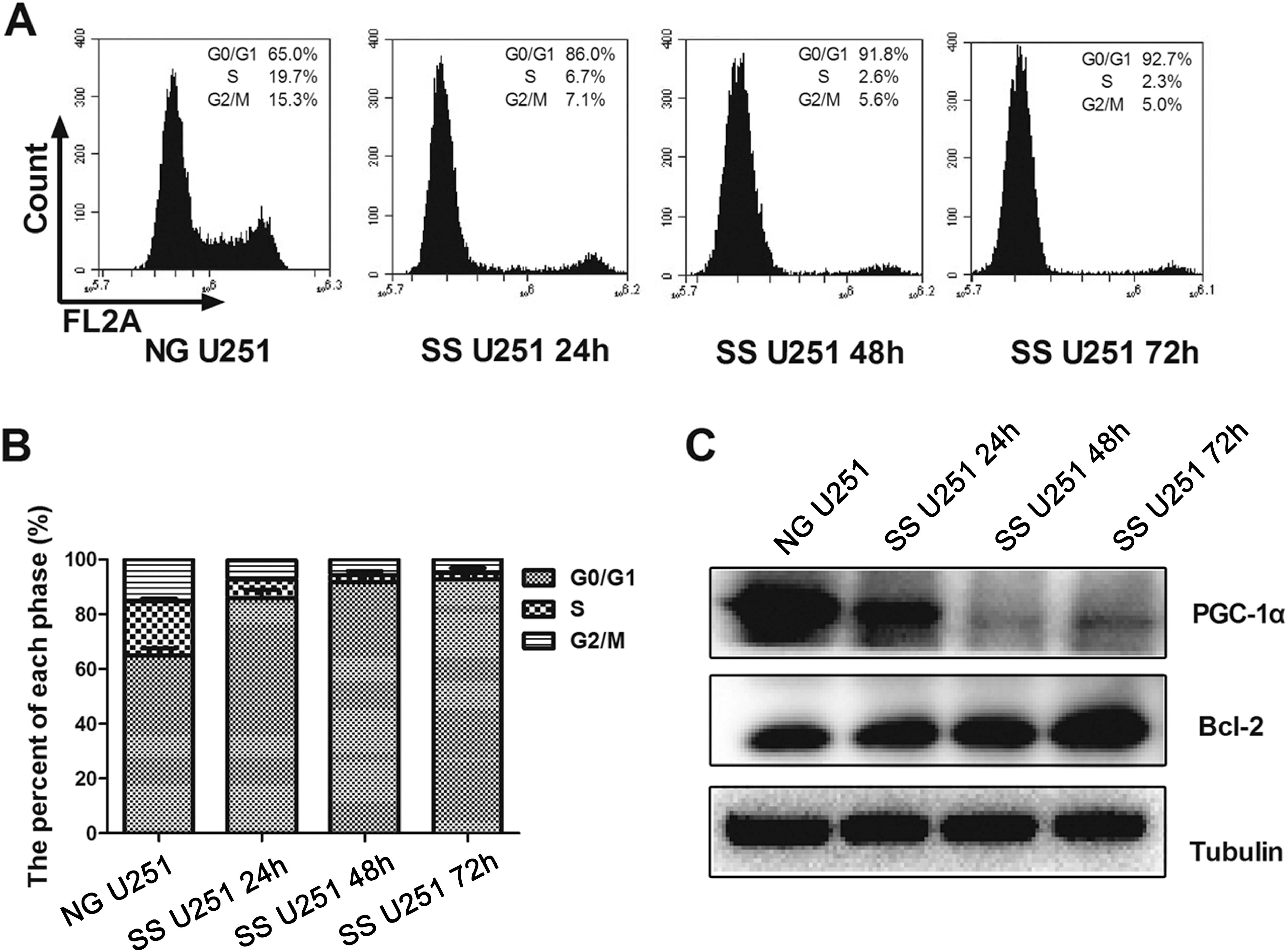
Cell cycle profiles and Bcl-2 and PGC-1α expression after SS treatment of U251 cells. (A) U251 cells were cultured in serum-free medium for 24, 48, and 72 h, and harvested for cell cycle analysis by flow cytometry. (B) The percentage of SS-treated or NG U251 cells in each cell cycle phase is shown. (C) PGC-1α and Bcl-2 expression in U251 cells after SS treatment was measured by western blot.

### Bcl-2 was upregulated following PGC-1α knockdown

To further test the relation between PGC-1α and Bcl-2, experiments with siRNA targeting PGC-1α were performed. Fig. 5A shows that PGC-1α was significantly decreased in cells transfected with siRNA, while no change was found in cells transfected with the negative control (NC), positive control (GAPDH) or the transfection reagent only (Fig. 5B), confirming the successful PGC-1α knockdown. Western blotting analysis showed that Bcl-2 expression was significantly increased after knockdown of PGC-1α (Fig. 5C). Based on these results and those shown in Figure 4, we conclude that Bcl-2 is negatively regulated by PGC-1α.

**Fig. 5.**
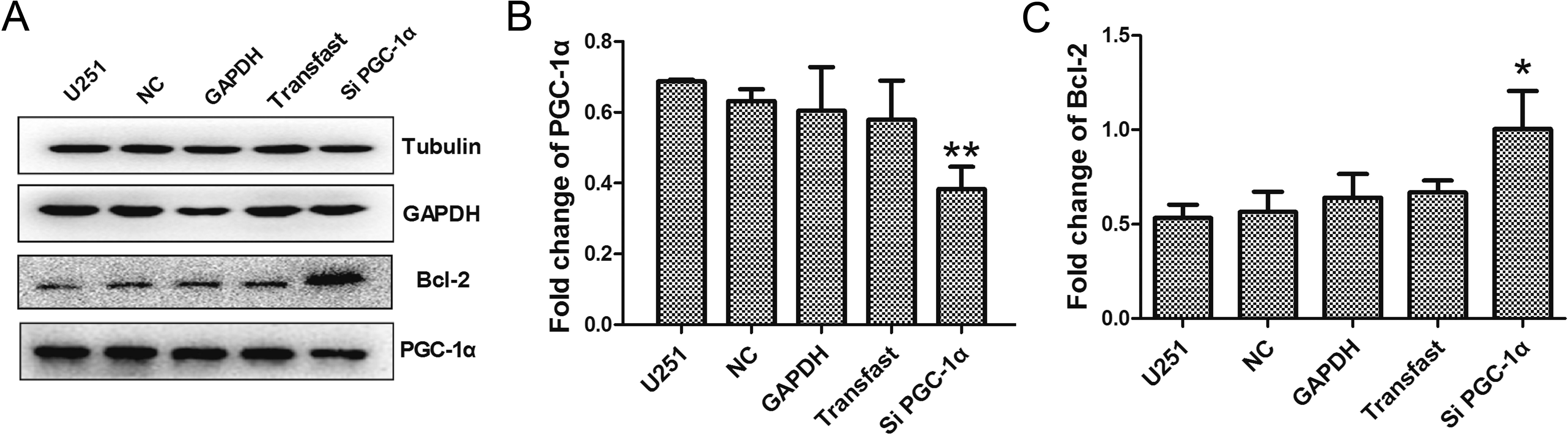
Bcl-2 was increased when PGC-la knockdown by PGC-la siRNA in U251 cells. (A) GAPDH, Bcl-2 and PGC-1α expression was measured by western blot after transfection with PGC-1α specific siRNA. (B) and (C) Quantification of Bcl-2 and PGC-1α protein expression shown in (A). NC: random siRNA serving as a negative control, GAPDH: positive control, Transfast: transfection reagents only.

## Discussion

The human Bcl-2 gene was first identified in tumor cells of follicular lymphoma patients, and was localized near the junction at which chromosomes 18 and 14 (t14;18) are joined (Tsujimoto et al., 1984). This chromosome translocation led to upregulation of Bcl-2 expression and contributed to cancer (Tsujimoto et al., 1985; Nunez et al., 1989). As an oncogene, Bcl-2 can inhibit apoptosis and promote cell survival, and these functions have been well characterized for decades. Bcl-2 binds to proapoptotic partners to form heterodimers that block cell death. The balance between anti- and pro-apoptotic functions of the Bcl-2 family proteins ultimately determines cell fate (Korsmeyer et al., 1993). However, the anti-tumor function of Bcl-2 is still not fully understood. Our group discovered that Bcl-2 could retard cell cycle progression through inhibiting p27 protein degradation (Janumyan et al., 2008; Du et al., 2017), but further investigation demonstrated that no direct interaction existed between Bcl-2 and p27 (data not shown).

Our recent work indicated that when the cells were synchronized at G_0_/G_1_ phase by SS, Bcl-2 overexpressing cells maintained a significantly lower ROS level compared to the control cells. However, there were no differences in ROS levels between normal growing Bcl-2 overexpressing cells and control cells. This is consistent with the fact that Bcl-2’s anti-tumor function was activated only in G_0_/G_1_ stage after serum starvation but not in normal growing status (Janumyan et al., 2008). Taken together, these results strongly indicate that ROS might play a role in Bcl-2’s cell cycle function (Du et al., 2017). Mitochondrial OXPHOS impinges on many cellular functions, including energy allocation and programmed cell death. ROS generated by mitochondrial OXPHOS provide a signaling system from mitochondria to the nucleus (Hansen et al., 2006), and are involved in cell cycle regulation, with low levels of ROS suppressing cellular growth and proliferation (Burdon et al., 1995; VanderHeiden et al., 2009). Bcl-2 was reported to have anti-oxidant effects as a ROS scavenger, but it lacks the sequence or structural features of known anti-oxidant proteins, suggesting that Bcl-2 might clear ROS by modulating mitochondrial bioenergetics (Nathan et al., 2009).

PGC-1α is a transcriptional co-activator of peroxisome proliferator-activated receptor gamma (PPARγ) that regulates transcription factor activity, and has a central place in multiple cellular processes, including mitochondrial respiration and OXPHOS (Alessia et al., 2010). PGC-1α responds to metabolic challenges such as exercise, starvation or cold; its expression is induced by these environmental stimuli and is associated with cancer (Yoon et al., 2001). We previously reported that as a master modulator of mitochondrial OXPHOS, PGC-1α reduced the ROS level to inhibit cell cycle (Fu et al., 2016). These findings led to the hypothesis that PGC-1α might directly or indirectly coordinate with Bcl-2 in regulating the cell cycle. To verify this hypothesis, we experimented with embryonic fibroblast NIH3T3 cells, in which Bcl-2 was either overexpressed or silenced, and which were arrested by SS or CI to activate the cell cycle function of Bcl-2. We observed that during serum starvation, the expression of p27 was increased significantly in G_0_/G_1_ arrested Bcl-2 overexpressing cells compared to control cells, which validated the cell cycle function of Bcl-2 (Janumyan et al., 2008; Janumyan et al., 2003; Cheng et al., 2004). PGC-1α was elevated as well in arrested Bcl-2 overexpressing cells but not in control cells (Fig. 1 and 2), suggesting a relationship between PGC-1α and Bcl-2. We hypothesized that Bcl-2 may recruit and rely on PGC-1α to reduce ROS, which mediated cell cycle as a critical signal. To test this possibility, Bcl-2 knockdown by siRNA was performed, and as we expected, PGC-1α expression was correspondingly decreased (Fig.3). Together with the facts that PGC-1α regulates mitochondrial function to eliminate ROS, and arrested Bcl-2 cells contain less ROS, these results suggest that Bcl-2 might recruit PGC-1α to reduce ROS levels and block cell cycle progression.

To further clarify the relationship between Bcl-2 and PGC-1α, we investigated the influence of PGC-1α knockdown on Bcl-2 expression. Endogenous expression of PGC-1α is low in normal growing 3T3Bcl-2 and control cells (Fig. 1C, Fig. 2C), and we failed to over-express PGC-1α in Bcl-2 cells for unknown reasons; thus human glioma U251 cells with high expression of endogenous PGC-1α were chosen to perform the experiments. Fig. 5 shows that Bcl-2 expression was upregulated after PGC-1α was successfully silenced. Consistent with this result, Bcl-2 was upregulated following PGC-1α downregulation during the SS treatment (Fig. 4). These results suggest a negative feedback between Bcl-2 and PGC-1α expression. Recently, the network between Bcl-2 and PGC-1α received intensive attentions due to their involvement in various types of cancers. In human musculoskeletal tumor cell lines, overexpression of PGC-1α increase mitochondrial numbers, which induce apoptosis (Onishi et al., 2014). Consistence with that work, Zhang et al. found that overexpression of PGC-1α in ovarian carcinoma cell lines significantly promotes apoptosis through reducing Bcl-2/BAX ratios (Zhang et al., 2007). Recent work reported that downregulation of PGC-1α decrease Bcl-2 expression and increase BAX expression that leads to apoptosis in human endometrial cancer cells (Yang et al., 2016). These findings suggested that PGC-1α play important roles through mitochondrial pathway with different patterns in various tumors. In present study, we elucidate the communication between PGC-1α and Bcl-2, mainly focusing on the anti-tumor function of Bcl-2 instead of its apoptosis role.

Elaborating on the possible mechanism of Bcl-2 on anti-tumor function is very important. Many studies have confirmed ATP and ROS were key signals in regulating cell cycle. Although it is known that Bcl-2 was involved in regulating cell cycle by ROS pathway, the mechanism of how Bcl-2 regulates the ROS pathway remains unclear. Bcl-2 lacks sequence or structural features of known anti-oxidant proteins (Nathan et al., 2009), suggesting that Bcl-2 indirectly controls ROS via other factors. PGC-1α was identified as a crucial switch on mitochondrial oxidative phosphorylation (Alessia et al., 2010), and function to diminish ROS (St-Pierre et al., 2006; Fu et al., 2016).

Based on our current study, we concluded that when Bcl-2 over-expressing cells (as most of the tumor cells) are confronting serum depletion, PGC-1α is activated, it might bind to Bcl-2 to inhibit cell cycle and suppress tumor cell proliferation through reducing ROS. Further study needs to be done to determine whether PGC-1α directly associate with Bcl-2 protein to control cell cycle. These findings revealed the communication between the mitochondria and the nuclear genes, provided the mechanistic understanding of Bcl-2’s anti-tumor function.

## Materials and methods

### Cell lines and cell culture

Human kidney epithelial 293T cells and human glioma U251 cells (with high Bcl-2 and PGC-1α endogenous expression) were supplied by the Biochemistry and Molecular Biology Laboratory of Yunnan University, P. R. China. Mouse embryonic fibroblasts NIH3T3 cells were purchased from the Shanghai Cell Bank of the Chinese Academy of Sciences. Cells were cultured in Dulbecco’s Modified Eagle Medium (DMEM) (Gibco, Logan, USA) supplemented with 10% fetal bovine serum (FBS) (BI, Logan, USA) and 1% penicillin/streptomycin (Hyclone, Logan, USA) at 37 °C in 5% CO_2_.

### Lentivirus virus packaging and stable cell transfection

Recombinant pBABEpuro-Bcl2 vector, packaging plasmid pCLECO, and X-tremeGENE HP DNA Transfection Reagent (Roche, Basel, Swiss) were added in DMEM, mixed gently, and incubated at room temperature for 20 min. The mixture was added drop-wise into 293T cells in a 10 cm plate. After 48 h, the supernatant containing pBABEpuro-Bcl-2 virus was collected and filtered through a 0.45-μm filter. NIH3T3 cells were then infected with the virus. After 48 h of culture, the aminoglycoside antibiotic puromycin (Gibco-BRL, Carlsbad, USA) was added into the medium at final concentration of 4 μg/mL to select NIH3T3 cells with stable expression of Bcl-2. Bcl-2 expression was confirmed by western blotting.

### Cell cycle synchronization and analysis by flow cytometry

For cell synchronization via SS, U251 or NIH3T3 cells were washed three times with phosphate-buffered saline (PBS) and cultured for 48 h in medium containing no FBS (U251 cells) or containing 0.1% calf serum (NIH3T3 cells). For cell synchronization via CI, the cells were allowed to reach confluence and then maintained in culture for 5 days. Synchronized cells by either method were harvested, washed with cold PBS, and incubated with a solution containing 50 μg/mL propidium iodide (PI) and 0.03% TritonX-100 at room temperature for 20 min. For each sample, at least 2 x 10^5^ cells/mL were analyzed with a BD Accuri C6 flow cytometer (BD Biosciences, San Jose, CA). Cell cycle profiles were calculated using the C6 software.

### Western blotting

Cells were harvested, washed twice with PBS, incubated in radioimmunoprecipitation assay lysis buffer (Beyotime, Jiangsu, China) on ice for 20 min, and centrifuged at 10,000 x *g* for 15 min at 4 °C. The supernatant was collected and protein concentration was quantified with a bicinchoninic acid (BCA) protein assay kit (Dingguo, Beijing, China). Supernatant samples (50 μg protein) were loaded onto a 12.5% polyacrylamide gel for sodium dodecyl sulfate-polyacrylamide gel electrophoresis (SDS-PAGE), and transferred to a polyvinylidene fluoride (PVDF) membrane at constant voltage (100 V) for 2 h. The membrane was then blocked with 5% milk and probed with primary antibody (at 1:1,000 dilution) overnight at 4 °C. After being washed 3 times with PBST (phosphate buffer saline with triton x-100), the membrane was incubated with a secondary antibody (1:2,000 dilution) at room temperature for 2 h, and the signal was developed with an enhanced chemiluminescence kit (Thermoscientific, Boston, USA). The quantification of relative protein expression based on western blot signals was performed using the ImageJ software. Antibodies against PGC-1α and p27 were purchased from Cell Signaling Technology Co. (CST, Boston, USA), anti-tubulin antibody from Beyotime Company (Jiangsu, China), and anti-Bcl2 antibody from Becton, Dickinson and Company (BD, USA). Light chain specific horse radish peroxidase (HRP) conjugated anti-rabbit IgG secondary antibody was purchased from Jackson Immunoresearch laboratories Inc. (Jackson, USA).

## Gene knockdown by small interfering RNA (siRNA)

siRNAs for human PGC-1α (sense: 5′-GUCGCAGUCACAACACUUATT-3′, antisense: 5′-UAAGUGUUGUGACUGCGACTT-3), control (sense:5′-UUCUCCGAACGUGUCACGUTT-3′, antisense:5′-ACGUGACACGUUCGGAGAATT-3′), and human Bcl-2, (Bcl-2-homo-506 sense: 5 -GGGAGAACAGGGUACGAUATT-3′, antisense: 5′-UAUCGUACCCUGUUCUCCCTT-3′, Bcl-2-homo-528 sense: 5′-GGGAGAUAGUGAUGAAGUATT-3′, antisense: 5′-UACUUCAUCACUAUCUCCCTT-3′, Bcl-2-homo-928 sense: 5′ -GAGGAUUGUGGCCUUCUUUTT-3′, antisense: 5′-AAAGAAGGCCACAAUCCUCTT-3′) were purchased from GenePharma. Transfections were performed according to the X-tremeGENE HP Transfection Reagent’s protocol. Briefly, 160 pmol siRNA and 10 μL X-tremeGENE HP Transfection Reagent were diluted in 50 μL DMEM respectively, incubated for 5 min, gently mixed together, and incubated for 15 min at room temperature to form siRNA-X-tremeGENE HP complexes, which were then added to 7 × 10^5^ cells per well plated in a 6-well dish and incubated for 48 h.

## Author contributions

Q. C. and X. D. designed the experiments and revised the manuscript. K. Y. and X. F. performed the experiments and wrote the manuscript text, Y. L., B. H., Z. C., and S. Y. prepared the experiments, R. W. and J. Z. analyzed the data.

